# Bacterial dynamin-like protein DynA mediates lipid and content mixing and shows phospholipid specificity

**DOI:** 10.1101/363689

**Authors:** Lijun Guo, Marc Bramkamp

## Abstract

The dynamins family of GTPases is involved in key cellular processes in eukaryotes, including vesicle trafficking and organelle division. The GTP hydrolysis cycle of dynamin translates to a conformational change in the protein structure, which forces the underlying lipid layer into an energetically unstable conformation that promotes membrane rearrangements. Many bacterial genomes encode dynamin-like proteins, but the biological function of these proteins has remained largely enigmatic. In recent years, our group has reported that the dynamin-like protein DynA from *Bacillus subtilis* mediates nucleotide-independent membrane tethering in vitro and contributes to the innate immunity of bacteria against membrane stress and phage infection. However, so far the mechanism of membrane stress response and the role of GTP hydrolysis remain unclear. Here, we employed content mixing and lipid mixing assays in reconstituted systems to study if the dynamin-like protein DynA from *B. subtilis* induces membrane full fusion, and further test the possibility that GTP hydrolysis of DynA may act on the fusion-through-hemifusion pathway. Our results based on fluorescence resonance energy transfer (FRET) indicated that DynA could induce aqueous content mixing even in absence of GTP. Moreover, DynA-induced membrane fusion in vitro is a thermo-promoted slow response. Surprisingly, digestion of protein mediated an instantl rise of content exchange, supporting the assumption that disassembly of DynA is the fundamental power for fusion-through-hemifusion.

## Introduction

Bacterial dynamin-like protein DynA in *Bacillus subtilis* belongs to the dynamin superfamily that includes classical dynamins in eukaryotes and dynamin-like proteins (DLPs). It is proposed that all dynamin-superfamily members are large mechanochemical GTPases involved in a variety of cellular processes and are major mediators of membrane remodeling. Structurally, dynamin family members are classified with large GTPase and coiled-coil region called middle domain (1,2). Most of the members share three properties: GTPase activity, oligomerisation and membrane binding.

Classical dynamin is the founding members of the dynamin family (3,4), functioning at the heart of endocytic vesicle fission in animal cells. Dynamins contain the following characteristic domains: GTPase domain, a helical stalk domain, pleckstrin homology domain (PH), and C-terminal proline/arginine rich domain (PRD). The PH domain mediates membrane binding and has specificity for phosphatidylinositol-4,5-bisphosphate. The stalks domain is involved in dynamin oligomerisation, and oligomerization is stimulating cooperative GTPase activity. Dynamins possess the remarkable property of assembling into contractile helical polymers that wrap around the neck of budding vesicles. Constriction of this helix contributes to severing the membrane and promotes vesicles release. The most common model is that the GTP hydrolysis cycle translates to a radical conformational change in the protein structure, which forces the underlying lipid layer into an energetically unstable conformation that promotes membrane rearrangements. However, there is still a possibility that membrane constriction is mediated merely by protein assembly (5).

A large family of dynamin-like proteins include mitochondria-associated OPA1/ Mgm1p-like proteins (6,7), Dnm1/DRP1-like proteins (8,9) and Mitofusins/Fzo1 (10,11), endocytosis-associated Vps1 like proteins (12), antiviral Mx proteins (13), anti-parasitic guanylate-binding proteins (GBPs) (14), ARC-5 like plant proteins (15) and bacterial DLPs (BDLPs) (16–19). These proteins catalyze a wide variety of membrane fission and fusion events, and some proteins also response to environmental stress. For example, Mx proteins in eukaryotic cells promote resistance not only to RNA viruses, but also to a wide range of DNA viruses, thus forming a part of the vertebrate innate immune system (20). The expression of GBPs is induced by type II interferon and these proteins have a role in resistance against intracellular pathogens (14), which is similar to, but less efficient than, that of the Mx proteins.

More than 900 bacterial species have been identified that contain DLPs. A BDLP from *Nostoc punctiforme* (*Nos*DLP), has been characterized as a membrane-binding GTPase, containing a GTPase, stalk domain and a paddle region identified by its crystal structure (16,21). *Nos*DLP contains a paddle domain that is supposed to allow for membrane binding, similar to the PH domain in dynamin. In *Bacillus subtilis*, dynamin-like protein DynA is head-to-tail fusion of two dynamin-like subunits and consequently possesses two separate GTPases (1,18). Either of two subunits shows high structural similarity with *Nos*DLP, except for the lack of the paddle region in D2 subunit. DynA catalyzes nucleotide-independent membrane binding and tethering in vivo and in vitro and helps cells to counteract membrane pore formation provoked by antibiotics and phages (2,18). The *Escherichia coli* LeoABC dynamin-like proteins play a role in potentiating virulence through membrane vesicle associated toxin secretion (22).

All BDLPs analyzed so far have a much lower GTP hydrolysis rate, compared to the eukaryotic DLPs, and purified LeoA does not show any GTPase activity and their role of GTP hydrolysis is not known (16–18). In order to uncover the mechanism how DynA works on membrane remodeling and responds to membrane stress, two fundamental questions need to be answered: i) are bacterial dynamin-like proteins alone sufficient to induce membrane full fusion (content mixing) and, ii) is GTP hydrolysisis involved in the step of membrane constriction, or protein disassembly, or neither. On a structural level, membrane full fusion results in the unification of the lipid and protein components and the intermixing of the volumes (23). Therefore, both lipid mixing and content mixing need to be tested for solving the above questions, and to directly verify the mixing of the content is the key. Several content mixing assays are already described for vesicle-fusion related SNARE proteins (24), such as fluorescence quenching assay with ANTS/DPX (25) utilizing the collisional quenching of polyanionic ANTS and cationic quencher DPX; fluorescence enhancement assay with Tb^3+^/DPA (26) were the chelate of Tb^3+^/DPA is 10,000 times more fluorescent than Tb^3+^ alone; fluorescence forming assay with vacuolar hydrolases/calcein-AM (27) based on that vacuolar hydrolases cleave the ester bonds of calcein-AM, yielding its bright green fluorescent form; single molecular DNA assay (28) based on the fact that a fusion pore allows two designed DNA molecules hybridize, opening up the stem region of the hairpin and then decreasing FRET efficiency between Cy3 and Cy5; and finally fluorescence FRET assay with Biotin-R-phycoerythrin(Bo-PhycoE)/streptavidin-Cy5(Sa-Cy5) (29), that exploits the high affinity between biotin and streptavidin. The later method of Bo-PhycoE/Sa-Cy5 is outstanding in terms of ease of use and responsiveness.

Here we employed content mixing assay of Bo-PhycoE/Sa-Cy5 and lipid mixing assays in reconstituted systems to study if the dynamin-like protein DynA from *Bacillus subtilis* could induce membrane full fusion and further test the possibility that GTP hydrolysis of DynA acts on the fusion-through-hemifusion pathway. We could show that DynA is able to catalyze lipid and content mixing in a GTPase independent manner. We further show that DynA preferentially binds to phospholipid mixtures that mimic the native membrane composition of *B. subtilis* cells.

## Results

Membrane fusion can be separated in various distinct steps: membrane tethering in trans, formation of hemifusion stalk (lipid mixing), and fusion pore expansion to the point at which the vesicle membrane flattens on the membrane interaction surface, leading to the release of the luminal contents (23). Membrane full fusion results in the unification of the lipid bilayer and the intermixing of the volumes. Hemifusion is the intermediate stage for membrane full
fusion that allows the exchange of lipids between the outer leaflets whereas lipid exchange between inner layers and content mixing is still blocked. However, docking of membranes is usually not sufficient for lipid exchange. To test DynA mediated membrane fusion, we employed assays based on Förster resonance energy transfer (FRET). Specifically, we used two lipid mixing assays, termed lipid FRET and lipid dequenching as described before (30). Furthermore an content mixing assay (termed content FRET here) was used to address whether DynA activity can lead to complete membrane fusion (29) (Figure 1). For lipid FRET assays, liposomes were prepared with fluorescent lipid Marina-Blue-1,2-Dihexadecanoyl-*sn*-Glycero-3-Phosphoethanolamme (MB-PE) or *N*-(7-Nitrobenz-2-Oxa-1,3-Diazol-4-yl)-1,2-Dihexadecanoyl-*sn*-Glycero-3-Phosphoethanolamine (NBD-PE), respectively. If the addition of protein can cause membrane vesicles to be semi-fused or fully fused, an increased FRET signal will be detected. For lipid dequenching assays, one set of vesicles is pre-formed with MB-PE and NBD-PE, while the other vesicles have no fluorescent lipids. Mixing with non-labeled vesicles and subsequent membrane fusion will quench the FRET signal. Bo-PhycoE and Sa-Cy5 were used as luminal reporters in content mixing assay. The labeled volume of the liposomes has the chance to mix and undergo FRET only when full fusion of the vesicles occurred. The high affinity of streptavidin and biotin allows the mixed content to generate an efficient FRET signal. For lipid dequenching and content FRET assays, the maximum values of reactions can be estimated by using the detergent thesit as control.

**Figure 1.**
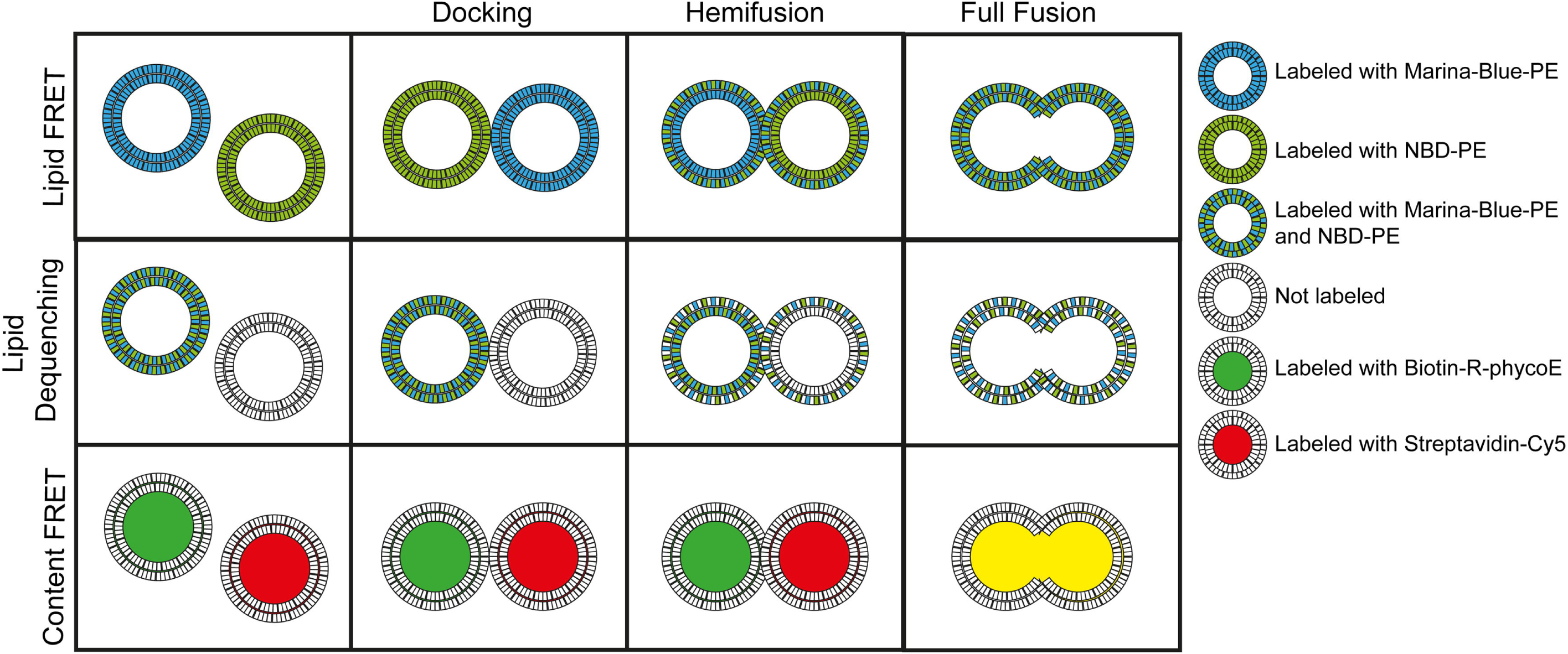
Cartoon of vesicle fusion assays used here to study DynA mediated membrane fusion.

### DynA induces nucleotide-independent lipid mixing leading to formation of large vesicles

DynA displayed nucleotide-independent membrane binding and tethering (18). Here, we further tested the ability of DynA in membrane fusion using the method of lipid FRET and observed the fusion process with fluorescence microscopy. The fluorescence signals of MB-PE and NBD-PE labeled vesicles showed no significant interference at the specific emission wavelengths (Figure 2Aa and 2Ab). When the two species of vesicles were mixed, the fluorescence signal was perfectly separated (Figure 2Ac) and, importantly membrane vesicles remain essentially intact ruling out that spontaneous fusion occurs in the time course of our experimental setup. After addition of DynA, membrane vesicles aggregated to form large membrane clusters (Figure 2Ad). These clusters revealed green and blue fluorescence. Addition of 1mM GTP gave essentially the same result (Figure 2Ae). In the protein-added samples, occasionally large, unilamellar vesicles (~ 8.918 μm) appeared (Figure 2Af, **Movie S1**), suggesting that DynA promoted the fusion of multiple small vesicles into large vesicles. In other words, DynA catalyzed full fusion of membrane vesicles in vitro. However, we did not observe any difference when GTP was added. We added Proteinase K to the protein-containing samples and incubated for ten minutes. Proteinase treatment lead to separation of membrane clusters into individual vesicles (Figure 2Ag **and S1A**). These vesicles were larger (~ 1.303 μm) than the vesicles in the sample without DynA that correspond to the size of the filter pores used for vesicle extrusion (~ 0.4 μm) and exhibited green and blue fluorescence. In order to get more quantitative data of the fusion event, we measured FRET efficiencies of lipid mixing assays (Figure 2B **and S1B**). The FRET data support the notion that there was no significant difference after addition of GTP (*P*= 0.43 for DynA:DynA/GTP, > 0.05; *P*=0.074 for DynA+PsK:DynA/GTP+PsK, >0.05) or after sudden digestion with proteinase K for 10min (*P*= 0.097, >0.05). Addition of GTP or proteinase did not promote the enhancement of lipid mixing efficiency, and it resulted in a slight decrease in fusion efficiency possibly due to changes in vesicle stability.

**Figure 2.**
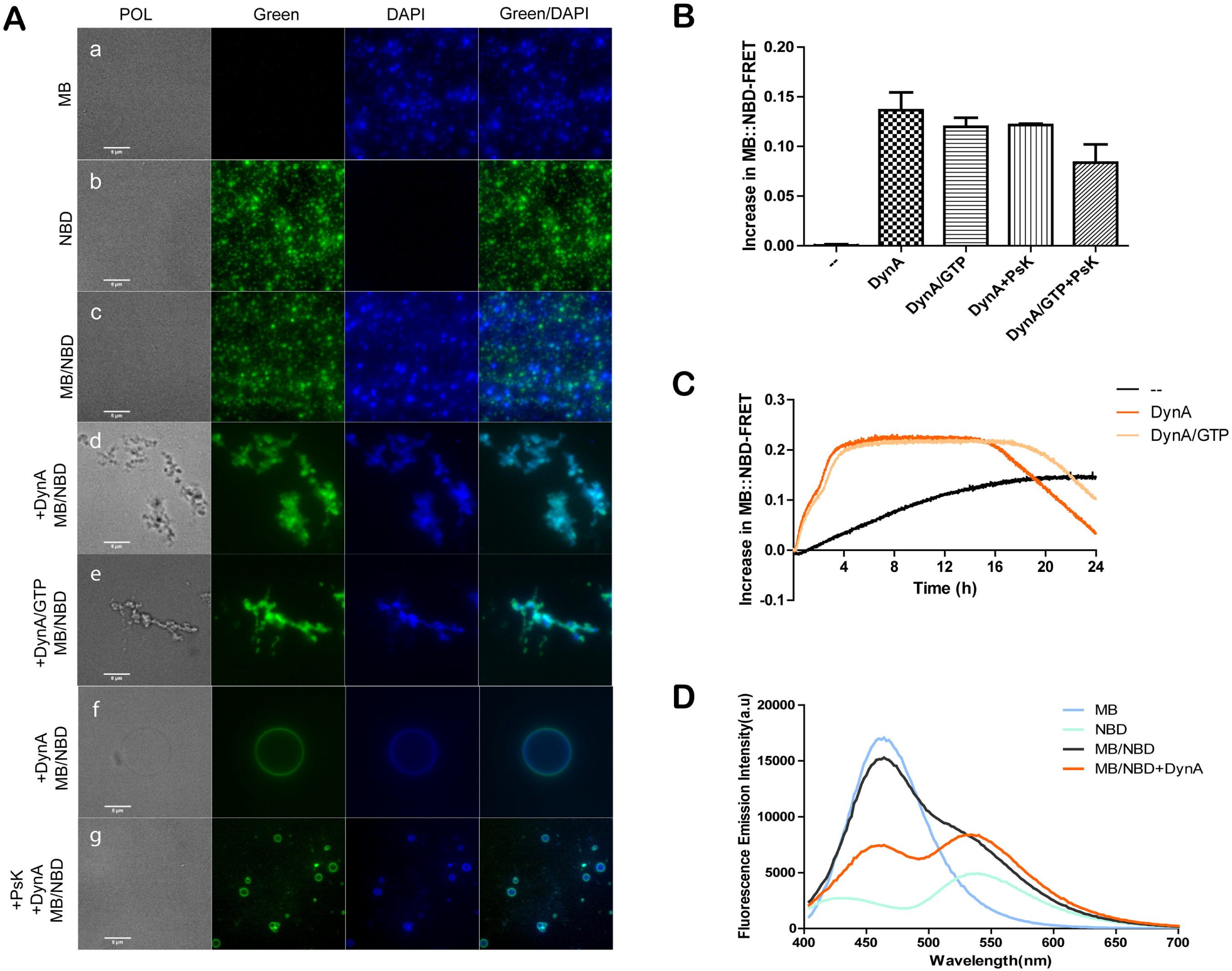
Nucleotide-independent lipid mixing induced by DynA. (**A**) Microscopic observation of DynA mediated vesicle fusion. (**a**) Vesicles labeled with MB (1 mol%) exhibited a blue fluorescence signal while (**b**) the NBD (1 mol%) labeled vesicles exhibited a green fluorescent signal. (**c**) After mixing the two species of vesicles (1:1) no fluorescent mixing was observed. (**d**) Purified protein DynA (0.5 μM) was added and incubated for 1h at 37°C. (**e**) Vesicle fusion in presence of 1 mM GTP. Note that in experiments with and without nucleotide large vesicle assemblies were observed. (**f**) Example of large unilamellar vesicles, a fusion product of DynA mediated membrane fusion. (**g**) After proteinase K treatment of the protein-containing samples, membrane clusters disengaged and larger sized membrane vesicles appeared. (**B**) Changes in Lipid-FRET efficiency after addition of DynA and proteinase K. The mean and SEM of the 5 replicates is shown. (**C**) Changes in FRET efficiency over a continuous 24 h period. (**D**) Fluorescence emission intensity at excitation of 370 nm with a ratio of MB and NBD vesicles of 1:9.

Next we wanted to address the time course of lipid mixing, therefore we monitored lipid-FRET efficiency over a long time period (0 - 24 h). Lipid FRET efficiency took approximately four hours to reach a stable plateau and started to decrease after 16h. The addition of GTP did not accelerate the fusion process. In control experiments without protein, the efficiency of fusion increased slightly because of a low degree of spontaneous fusion events. By increasing the amount of NBD-labeled vesicles, FRET efficiency can be significantly improved (**Figure S1E**). Optimization of FRET efficiencies revealed that a ninefold excess of FRET receptor (NBD-PE) over MB-PE clearly increased of MB::NBD-FRET efficiencies (Figure 2D **and S1B**). Additionally, protein concentrations directly affected the efficiency of membrane mixing. We determined optimal protein concentration to be 0.5 μM (**Figure S1C**). Since we did not observe any effect of GTP on membrane fusion, we wanted to test the possibility that a GTP effect may be masked by an overall high DynA concentration. In order to test this possibility, we used lower protein concentrations (100 nM, 50 nM and 10 nM) and found that adding GTP in all cases did not improve the efficiency of membrane fusion (**Figure S1D**). We have used different concentrations of GTP, because a large excess of nucleotide compared to protein concentration has inhibitory effects on the overall activity. At higher concentrations, GTP significantly reduced membrane fusion efficiency. In summary these data confirm that DynA mediates nucleotide independent lipid mixing.

### DynA can induce aqueous content mixing, even in absence of GTP

One limitation of the lipid mixing assays is that it does not directly discriminate between stages of hemifusion and full fusion. Although, we had successfully observed large vesicles caused by DynA, which suggested that DynA induced full membrane fusion, we also need to directly test the exchange of content to further confirm the suggestion and to provide quantitative data. Therefore, we used content FRET to further explore the features of membrane fusion induced by DynA. First, we optimized the preparations to minimize fluorophore leakage. For this purpose, we directly checked the membrane vesicles by fluorescent microscopy and found that the donor or the receptor vesicles could be observed under the orange or Cy5 channel, and the fluorescence signals of these two did not interfere (Figure 3Aa and 3Ab). After mixing, their fluorescence signals did not overlap, suggesting that spontaneous content mixing occurs if at all at low background levels (Figure 3Ac). After addition of DynA, vesicles aggregated, as seen before. Unlike in lipid mixing assay, we did not observe immediate mixing of the fluorescent dyes in the content mixing assays, but were able to see single labeled vesicles in the tethered vesicle clusters (Figure 3Ad). Addition of GTP also did not cause differences in vesicle clustering as observed microscopically (Figure 3Ae). Occasionally, large vesicles appeared that exhibited red and green fluorescence (Figure 3Af).

**Figure 3.**
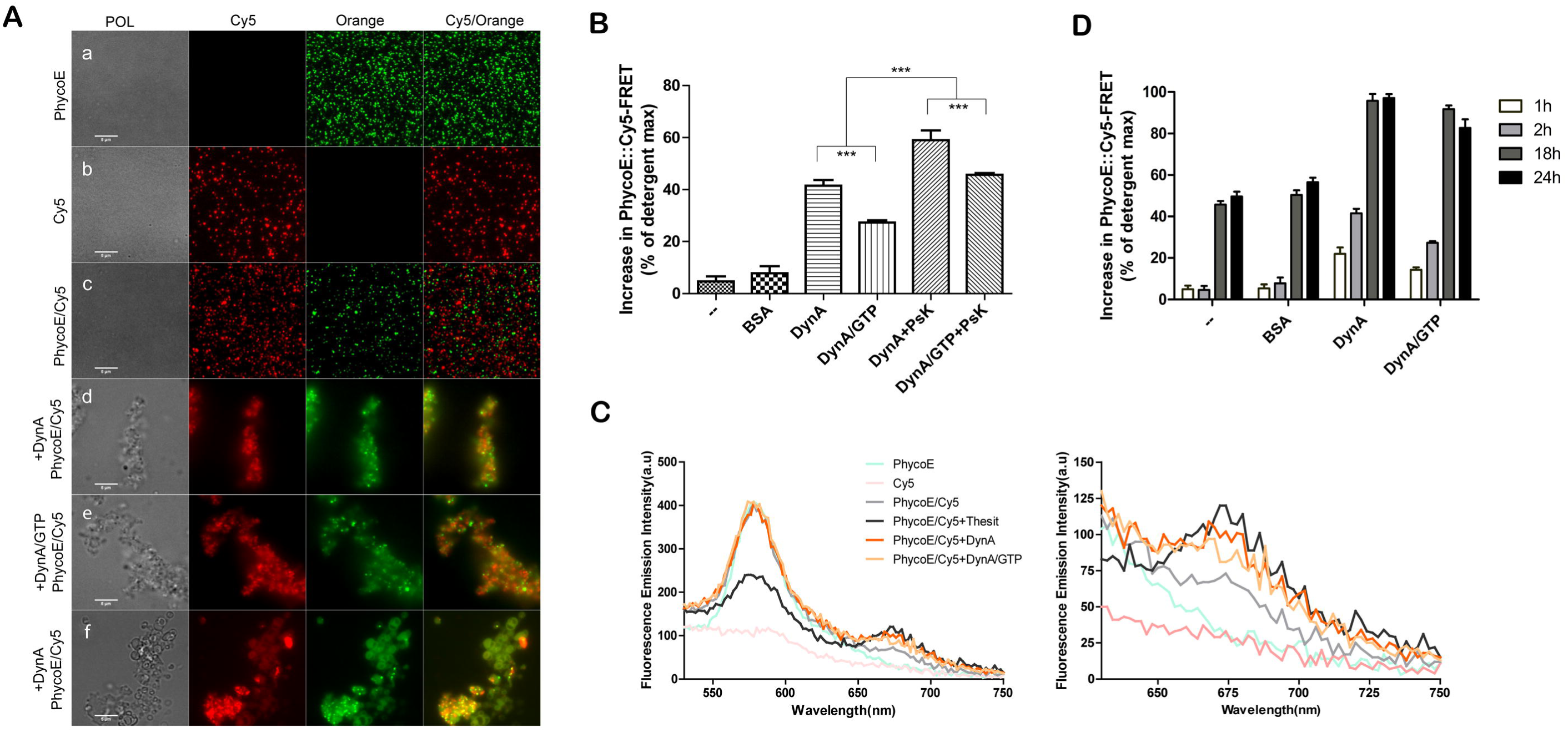
DynA induces aqueous content mixing. (**A**) Microscopic observation of DynA mediated content mixing. (**a**) Vesicles were labeled with Bo-PhycoE (0.4 μM) in their lumen or (**b**) loaded with Sa-Cy5 (0.4 μM). (**c**) Mixing of the two differently labeled vesicles did not result in fluorescent overlapping, indicating that vesicles stayed intact and did not fuse spontaneously. (**d**, **e**) After addition of DynA (0.5μM) and incubation for 1h at 37°C with or without 1mM GTP, large membrane clusters were formed. (**f**) In the case of longer reaction time (16h), larger vesicles were observed. (**B**) Changes in content-FRET efficiency after addition of DynA and proteinase K. Error bars are SEM of 5 replicates. (**C**) Fluorescence emission intensity after excitation at 496 nm with a 1:1 ratio of Bo-PhycoE and Sa-Cy5 inside the vesicle lumen. (**D**) Changes in FRET efficiency at various reaction periods (1 h, 2 h, 16 h and 24 h).

Addition of DynA led to an increase in content FRET efficiency, directly indicating that DynA-induced membrane full fusion (Figure 3B and 3C). We found that the short-term treatment with Proteinase K could lead to an increase in the proportion composed by total fusion (*P*=0.00021, < 0.05). This showed that DynA detachment from the membrane surface is an essential step for the transition from hemifusion to full fusion. As the result of lipid FRET assay, GTP did not promote the increase in FRET efficiency, and content FRET could increase continuously over a long period (Figure 3D). Notably, GTP hydrolysis fails to promote lipid mixing, nor act on fusion-through-hemifusion pathway. Membrane full fusion can be induced slowly by DynA alone.

In addition, we simultaneously applied content mixing and lipid dequenching, that is, labeling vesicles membranes and content at the same time. The donor vesicles were labeled with MB-PE and NBD-PE, and contained Bo-PhycoE within their lumen, while the receptor vesicles did not contain FRET-label in the membrane, but contained Sa-Cy5 in the lumen. Fluorescence microscopy inspection revealed that the donor vesicles had fluorescent signals in three channels, green, blue, and orange as expected (Figure 4Aa), while the receptors had fluorescent signals only of Cy5 (Figure 4Ab), indicating that the prepared membrane vesicles met the test requirements. After addition of DynA, vesicles aggregated into clusters with fluorescent signals in all four channels (Fig. 4Ad). Within a span of 24 hours, content mixing reached a maximum at approximately 3.5 h (Figure 4C), and lipid dequenching reached a maximum at approximately 1.5 h (Figure 4B), suggesting DynA-induced content mixing lagged behind the lipid mixing. This two-hour delay shows the existence and stability of the intermediate state of hemifusion. The detergent maximum value of content mixing is much larger than that of lipid mixing, indicating that lipid mixing is faster and more efficiently catalyzed by DynA compared to content mixing. Larger PhycoE/MB/NBD vesicles (Figure 4Ab and 4Ac) were observed that exhibited stronger lipid fluorescence signals and had weaker content signals (white arrows). We believe that the bigger the vesicle, the weaker the content fluorescence intensity, due to leakage and vesicle instability. This can also explain the poor fluorescence of the inner lining of large vesicle in Figure 3Af.

**Figure 4.**
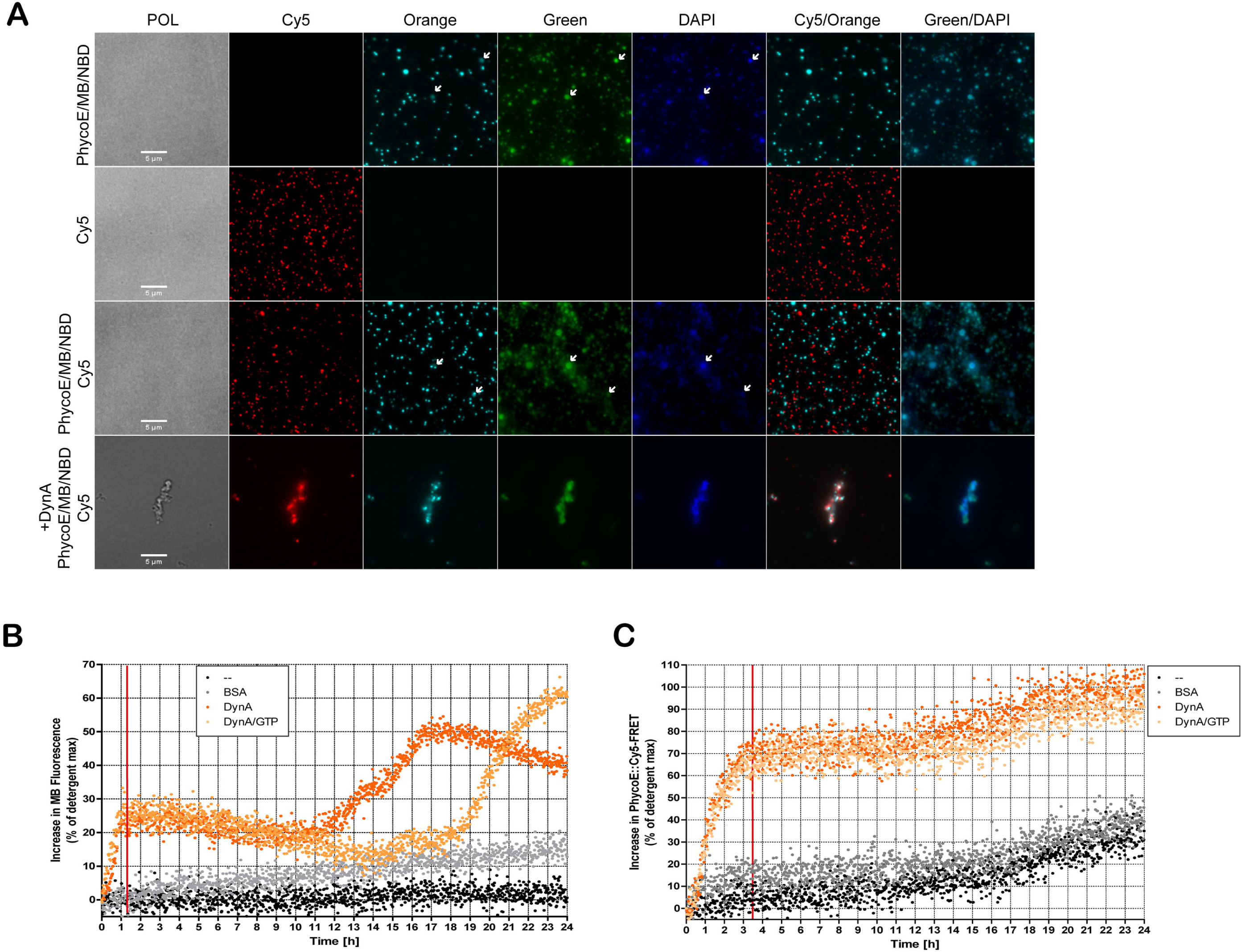
Combination analysis of membrane dequenching and content mixing. (**A**) Microscopic analysis of DynA effects on membrane and content labeled vesicles. (**B**) Changes in lipid dequenching efficiency over a continuous 24 h period. Lipid dequenching efficiency was shown by the increase of MB fluorescence. Shown is the mean of the 5 replicates. (**C**) Changes in content FRET efficiency over a continuous 24-h period. Shown is the mean of the 5 replicates.

### Membrane fusion is thermally accelerated

We found that at ambient temperature, 24 °C, membrane fusion was lower compared to 37 °C (**Figure S2A**). We reasoned that membrane fusion is likely influenced by temperature. Therefore, we compared DynA-mediated fusion in detail at 24 °C and 37 °C. When we raised the temperature to 37°C, the number of large vesicles observed microscopically was increased. Since lipid mixing was clearly enhanced at 37 °C compared to 24 °C, we assumed that 37 °C would be a more suitable temperature to analyze content mixing. Content FRET could be easily detected at 37 °C (**Figure S2B**), indicating that higher temperatures have a positive effect on full membrane fusion and content mixing. Similarly, we measured the FRET efficiency over time by incubating samples at 24°C and 37°C. Membrane fusion efficiency reached its maximum in less than 2 h at 37 ° C, while it took more than 4 h at 24 °C (Figure 5). Moreover, the fusion efficiency at 24°C was always lower compared to that detected for incubation at 37°C. Also, an increase in temperature increased FRET background over a 24 h period, indicating a higher degree of spontaneous fusion events.

**Figure 5.**
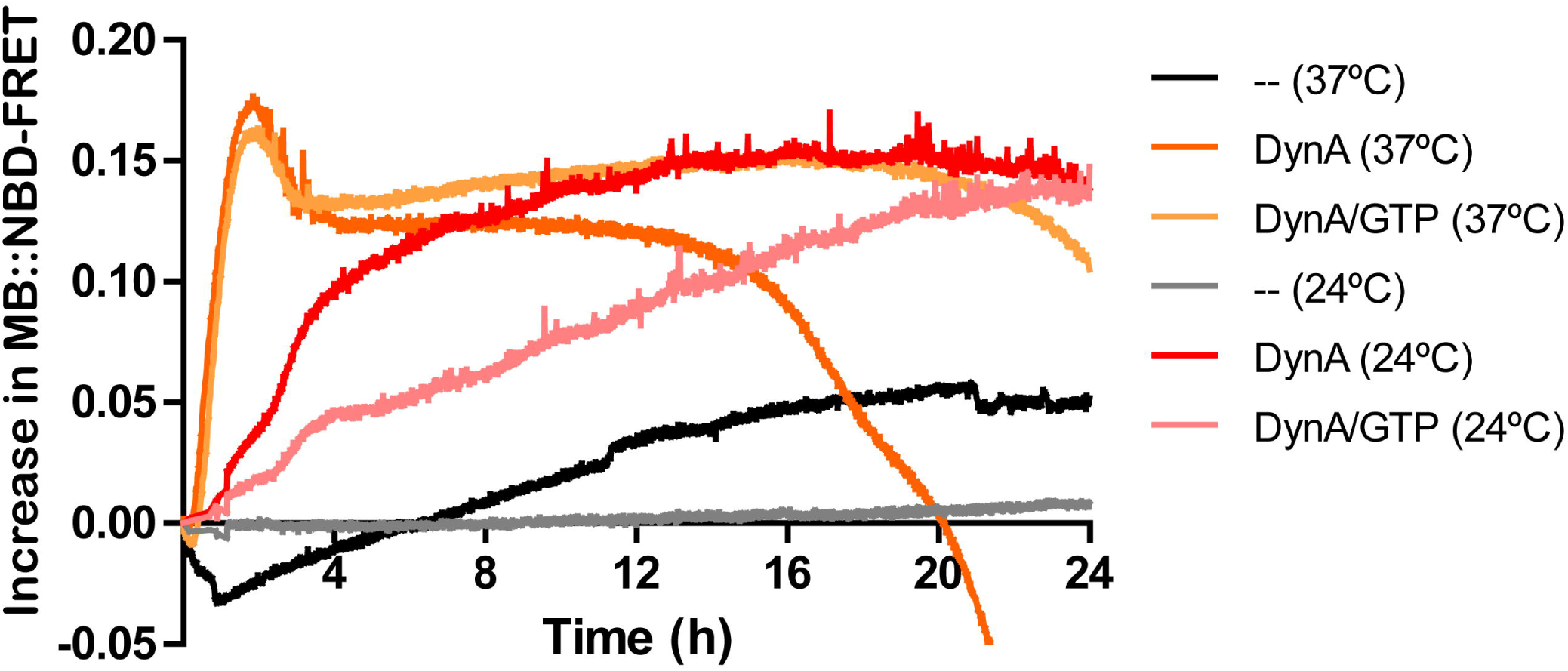
Thermal effects on DynA mediated membrane fusion. Changes in lipid-FRET efficiency were measured over a continuous 24 h period at 24 °C and 37 °C. The mean of the 5 replicates is shown. Ratio between MB and NBD vesicles was 1:9.

### DynA exhibits phospholipid preferences

Many lipid-binding proteins associate with specific lipids (31). To test whether DynA exhibits lipid specificity, we designed synthetic lipid vesicles with varying amounts of different phospholipids. The lipid for membrane fusion assays used in previous experiments was extracted from *E. coli* and contained 57.5 % (wt/wt) phosphatidylethanolamine (PE), 15.1 % phosphatidylglycerol (PG) and 9.8 % cardiolipin (CL). *B. subtilis* contained at least five major phospholipids, four of which were isolated and identified as a polyglycerol phospholipid, specifically PE, PG, CL, and lysylphosphatidylglycerol accounting 36 %, 30 %, 12 % and 22 % when cells are grown in rich medium (32). For *E. coli* and *B. subtilis*, PE, PG and CL are important phospholipid components, in which PG and CL are anionic membrane phospholipids. PG acts to maintain bacterial cell membrane stability (33). CL is a unique phospholipid with dimeric structure that carries four acyl groups and two negative charges whose structure changes in the presence of pH and divalent cations. Additionally, CL is believed to play a role in membrane fusion (34). PE, in addition to its structural role in membranes serves multiple important cellular functions including mitochondrial fusion in eukaryotes (35–37).

To test the binding preference of DynA for the phospholipids, we reconstructed membrane vesicles with different ratios of phospholipids in vitro and performed a lipid dequeching assay. Similar to lipid FRET assay, a higher ratio of acceptors (NBD-PE) to donors (MB-PE) resulted in more efficient FRET dequenching (**Figure S3**). Thus, in the experiment, the ratio of FRET donor to of FRET receptor was 9:1. When vesicles were prepared only with PE membrane fusion was not detectable (Figure 6A). However, with an increase in PG ratios, membrane fusion efficiency increased significantly, reaching the maximum at PG=40% and then gradually decreased. Membrane fusion could also be observed when vesicles contained only PG, indicating that PG is required and sufficient to recruit DynA to the membrane. In order to test the influence of CL, we constructed vesicles with a radio of PG to PE as 2:3 and added different amount of CL, and found that when CL was around 30 %~40 % the fusion efficiency could be improved (Figure 6B).

**Figure 6.**
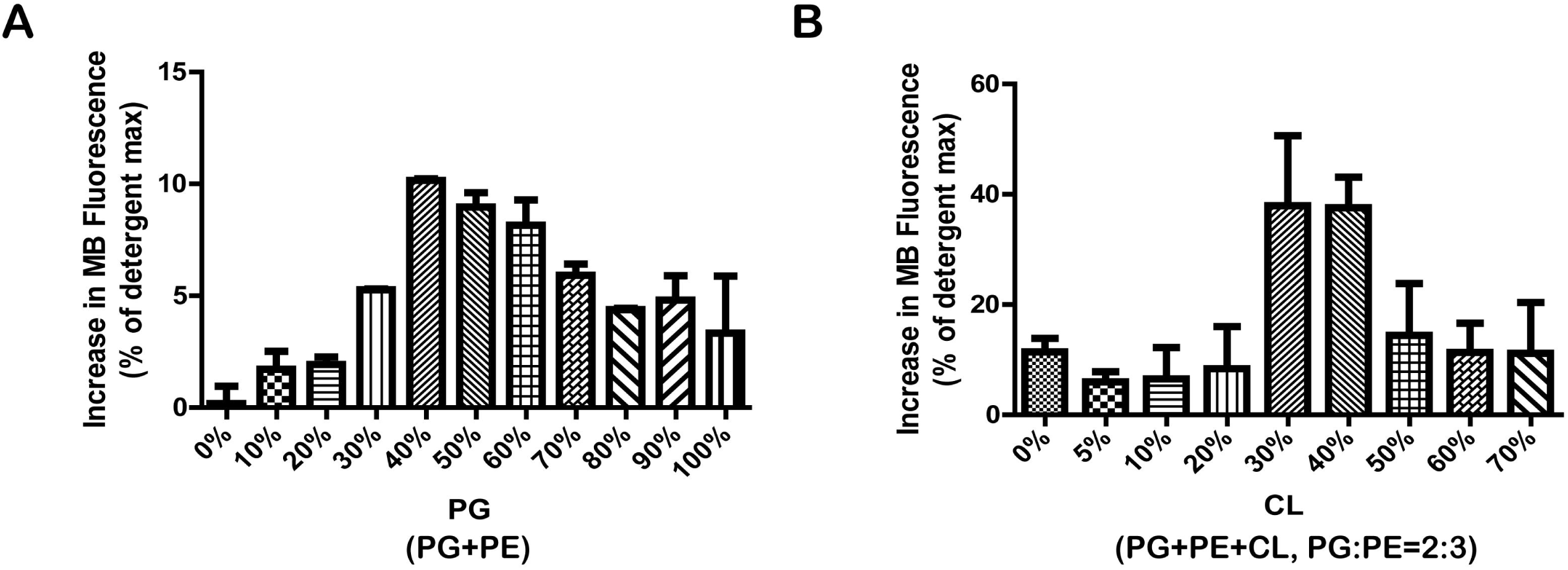
DynA shows phospholipid preferences. Vesicles were extruded from PG, PE, and CL, and labeled for lipid dequenching assays. Vesicles were incubated with DynA at 24 °C for 30 min. (**A**) DynA mediated lipid mixing is lipid specific and requires PG. (**B**) CL at concentrations of 30-40 %, promoted lipid mixing. Mean and SEM of 3 replicates are shown.

### The D1 subunit of DynA is crucial for membrane fusion

Bacterial dynamin-like protein *Nos*DLP homo-dimerizes in its GDP-bound state via its GTPase domain, and in the presence of GTP and lipids, the protein self assembles around the liposome and forms a lipid tube (16). However, we have shown before that most bacterial dynamin-like proteins may act as heterooligomers since they are usually encoded as two copies in an operon (as in *N. punctifome*) or the two genes are fused in a head-to-tail fashion giving rise to a fusion protein with two dynamin-like subunits (18). *B. subtilis* DynA is such an internal fusion protein containing two BDLP subunits (38). Earlier work showed that membrane binding is achieved via the D1 part of DynA (18). Here we tested the membrane fusion efficiency of these two subunits and the full-length protein. Compared to full-length protein and D1 subunit, membrane fusion induced by D2 subunit was much less efficient (Figure 7A and 7B), indicating that D1 is the main driving force of membrane fusion. A complementation in trans with soluble D1 and D2 parts of DynA does not reconstitute membrane fusion, indicating that a tight interaction and coordination between the two parts of DynA is essential (data not shown). These results show that D1 and D2 play different roles in the process of membrane fusion.

**Figure 7.**
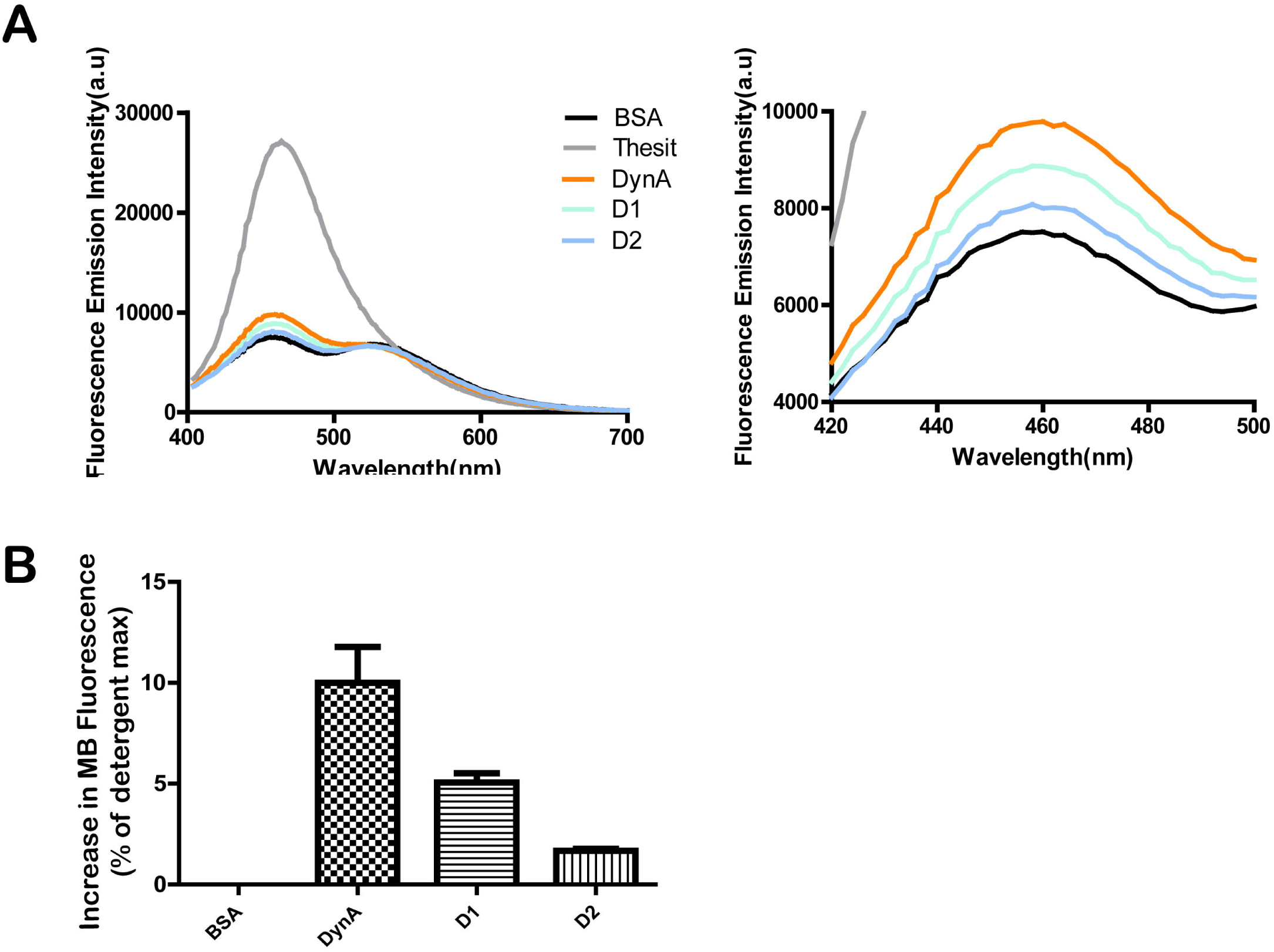
Isolated D1 and D2 subunits of DynA have reduced activity. Comparison between full length DynA and the D1 and D2 subunit in lipid dequenching assays. (**A**) Fluorescence emission intensity after excitation at 370 nm. (**B**) Lipid dequenching efficiencies after addition of protein. Mean and SEM of 3 replicates are shown.

## Discussion

The *B. subtilis* DLP DynA is a head-to-tail fusion of two DLP subunits and was shown to dimerize in vitro (18). We have shown earlier with in vitro and in vivo experiments that DynA is able to tether membranes in trans and is able to promote lipid mixing (18). Evidence was provided that DynA plays a protective role when cells were challenged with membrane pore forming agents such as the antibiotic nisin or cells were infected with phages (2). Based on these phenotypes it was suggested that DynA may seal effectively membrane pores. Similarly, in *Streptomyces* it has been proposed that DLPs are involved in membrane fusion during cytokinesis (19). These cellular roles would clearly require fusion of both membrane leaflets and hemifusion would not be sufficient to repair damaged membranes. Therefore, we set out to investigate the fusion activity of DynA in detail with a series of in vitro fusion assays. Using FRET based fusion assays we provide evidence here that DynA is indeed able to promote full fusion of membranes. The D1 part of the enzyme shows higher affinity for membranes compared to D2, but optimal activity was only achieved with the full length protein. A mixture of D1 and D2 does not restore effective fusion activity, indicating that a tight coupling of the two DynA parts exists and that this coupling is required for function. Tethering and fusion is in vitro apparently independent of nucleotide hydrolysis, provoking the question why these enzymes have a measurable GTP hydrolysis activity. Our in vitro fusion assays reveal that full membrane fusion is a slow process in vitro. This slow membrane fusion is not compatible with the role of DynA in membrane protection after pore formation. In vivo, the membrane potential would collapse when membrane pores are not quickly and effectively sealed. Interestingly, we observed that removal of DynA via proteinase K treatment, leads to a rapid increase in fusion activity. We assume that DynA assembly on opposing membranes leads to a membrane deformation that is energetically unfavorable and removal of membrane bound DynA would allow efficient membrane fusion. Thus, GTP hydrolysis might be involved in the rapid removal of DynA from the membrane. This is difficult to test in vitro and even experiments with very low concentrations of DynA (10 nM) did not reveal any positive effect of GTP on fusion activity. It may therefore be likely, that we still lack a crucial cellular factor that would speed up membrane fusion in vivo and that would require nucleotide turnover.

It is suggested that the first step of fusion-through-hemifusion is fusion pore opening and this step is limited by a larger free energy barrier than the induction of hemifusion (23,39,40). This would be in line with our hypothesis that GTP hydrolysis provides energy for the jump process from hemifusion to full fusion. Since the beginning of the lipid mixing is a logarithmic increase, the transition from docking to hemifusion is instantaneous. After the vesicles in the cluster complete hemifusion, they begin to gradually transition to full fusion. That is, the outer layer is flattened on the inner layer, and the inner layer begins to exchange membrane components. After that, the fusion pores begin to form and subsequently enlarge, followed by content exchange through the enlarged fusion pore, resulting in appearance of large vesicle. Since the time for the content mixing to reach the maximum is two hours later than the lipid mixing, we believe that the formation and expansion of the fusion pore is a slow process at least under the observed in vitro situation. Next to the gene encoding DynA two small OPFs are present (*ypbQ* and *ypbS*). We have isolated YpbS and checked for any influence on the in vitro fusion, but did not detect and difference (data not shown). YpbQ purification was not successful. Thus, it remains unclear how and why GTP turnover is an intrinsic property of DynA. Membrane fusion is in general a tightly controlled pathway. Fusion of synaptic vesicles with the plasma membrane is catalyzed by soluble N-ethyl maleimide sensitive factor attachment protein receptors (SNAREs) (41). SNARE mediated membrane fusion shares some similarities with the DLP mediated membrane fusion. First, v-SNAREs and t-SNAREs tether vesicle and plasma membrane and lead to entropy driven membrane tethering and fusion (42). It is assumed that the transmembrane helices of SNAREs play an important role in regulation of the fusion pore construction. Since bacterial DLPs such as BDLP1 (16), LeoA (17) and DynA (18) lack a clear transmembrane helix and rather insert with a paddle domain in one leaflet of the membrane, one might speculate that a so far unknown membrane protein could be involved in bacterial membrane pore fusion. In SNARE mediated membrane fusion several proteins are involved at various steps of membrane fusion (41). It seems plausible that also DynA may not act alone on the membrane. SNARE mediated vacuole membrane fusion is coupled to nucleotide hydrolysis in the Sec17 and Sec18 proteins. Interestingly, Sec17 and Sec18 act nucleotide hydrolysis independent in membrane fusion (43), but require ATP hydrolysis for the disassembly of the SNARE complex (44). Sec17 becomes essential for fusion when conditions such as membrane composition or SNARE complex assembly are not ideal (45). Thus, a certain mechanistic similarity between the SNARE mediated membrane fusion and the function of the bacterial DLP DynA is apparent. Several dynamin-like proteins show oligomerization and promote homotypic membrane fusion. The outer mitochondrial membrane is fused by the DPLs MFN1/2 in mammals (46) and FZO1/2 in yeast (47) and inner membrane fusion is triggered by optic atrophy 1 (OPA1) (48) or Mgm1 (49). The endoplasmic reticulum is fused by atlastin (50–52). These proteins all share a bacterial DLP-like molecular architecture, supporting the notion that DynA and other bacterial dynamins are also membrane fusion catalysts. Studies with MFN revealed that the two heptat repeat motifs (HR1 and HR2) are required for mitofusion function and HR1 likely perturbs the lipid bilayer, thereby promoting membrane fusion. Thus, DLP mediated membrane fusion may be triggered by local membrane perturbation rather than bridging complex formation as in SNARE complexes (53). HR1 was shown to bind to membrane directly. Due to its amphipathic helix structure, HR1 perturbs lipid order, rendering the deformed membrane fusogenic. DynA is supposed to insert with a paddle domain in the D1 part of the protein and not traverse the entire membrane with a transmembrane helix. Without detailed structural data on DynA it remains unclear whether the central part of DynA may be functionally homologous to the HR1 domain of mitofusin. Despite these molecular differences, SNARE and DLP mediated fusion have in common the deformation of membrane and the protein mediated trans tethering that is required for membrane fusion.

DynA binds best to membranes composed of a mixture of PG and PE in ratios similar to the situation in *B. subtilis* membranes. The cone shaped lipid PE has also been suggested essential for mitochondrial fusion (53) since it is enriched at mitochondrial contact sites (37). We also found that cardiolipin was able to increase the fusion efficiency, but only at high concentrations between 30-40%. In vivo CL concentration within the cell membrane are lower, however, we cannot rule out that CL may enrich in deformed membrane areas and that this local increase may have a positive effect on membrane fusion. In general, lipids with smaller head-groups seem to have a positive effect of DLP mediated membrane fusion. Physiological membrane composition has also been shown to be important for functional reconstitution of SNARE mediated membrane fusion (30,54,55). We also compared the effect of changes in temperature on membrane fusion, and found that the conditions closer to the optimum growth temperature of *B. subtilis* allow higher membrane fusion efficiency. Similar results were obtained for SNAREs that higher ambient temperature can promote membrane fusion (56,57). Likely, the increased dynamics of the phospholipids render the membrane at 37°C more fluid than at 24°C and thus allow faster membrane fusion to occur.

In summary, we have shown that the *B. subtilis* DLP DynA mediates full membrane fusion and that specific lipids such as PG are required to effectively exert its function. The data are in line with a role of DynA in membrane surveillance and protection against pore forming agents.

## Experimental Procedures

### Protein Purification

The *dynA* gene was cloned from *B.subtilis* 168 and inserted into pET16b(Novagen) with a C-teminal His_6_ tag. Expression was performed overnight at 18 °C with *E.coli* BL21(DE3) in Lysogeny broth (100 μg ml^−1^ carbenicillin) with 0.7mM IPTG. Cells were disrupted in 50 mM Tris, 200 mM NaCl, 20 mM imidazole, 10 % glycerol, 0.7 % TritonX-100, pH 8.0, and the protein was bound to Ni-NTA agarose (Qiagen). After extensive washing with 50 mM Tris, 200 mM NaCl, 200 mM imidazole, 10 % glycerol, pH 8.0 protein was eluted in 50 mM Tris, 500 mM NaCl, 1 M imidazole, 10 % glycerol, pH 8.0, reduced with 1mM DTT and gel filtrated on Superose 6 (GE Healthcare) against 50 mM Tris, 500 mM NaCl, 10 % glycerol, pH 8.0. Purification of DynA_K56A/K625A_, DynA subunit D1 (residues 1-609) and D2 (residues 561-1193) was analogous, except that lysis buffer of D1 subunit contained 500 mM NaCl.

### Liposome Preparation

*E. coli* total lipids were obtained from Avanti Polar Lipids, synthetic phospholipids of phosphatidylethanolamine(PE), phosphatidylglycerol (PG) and cardiolipin (CL) were from Sigma-Aldrich, and fluorescent lipids (Marina-Blue-PE and NBD-PE) were from Thermo Fisher Scientific. Following mixing lipid stock solution, lipid films were evaporated under a stream of nitrogen and further dried completely under vaccum. For liposome formation, lipids were diluted to 1 mg ml^−1^ (contains 1 mol% fluorescent lipids for lipid mixing assays) in RB150/Mg^2+^ (20 mM HEPES, 150 mM NaCl, 5 mM MgCl_2_, 10 % glycerol, pH 7.4), vortexed vigorously to get a homogenous solution and then extruded 20 times through a Millipore filter of pore size 0.4 μm. Liposomes were used directly after preparation, or stored at −80 °C. For content mixing assays, lipid concentration was 2 mg ml^−1^ in RB150/Mg^2+^ with 0.1% NaN_3_. Additionally, either of two luminal markers, Bo-PhycoE (Thermo Fisher Scientific, 0.4 μM) or Sa-Cy5 (Sigma-Aldrich, 0.4 μM) was added in lipid solutions before extrusion. Solutions were extruded 30 times and then dialyzed in 1000 kDa tubing (Spectrumlabs) for the Bo-PhycoE vesicles and 300 kDa tubing (Spectrumlabs) for the Sa-Cy5 vesicles three times (6 h, 12 h, 12 h) against 1000-fold volumes of RB150/Mg^2+^+ NaN_3_ to remove non-entrapped probes. In order to improve liposome quality excess Biotin was removed by using streptavidin magnetic beads (Sigma-Aldrich) and the incubated 30 min. Similarly, the streptavidin labeled vesicles were further cleaned with biotin magnetic beads from Raybiotech. For combination analysis of lipid dequenching and content mixing, lipid concentration was 2 mg ml^−1^ with 0.1 mol% fluorescent lipids and the luminal markers were 0.4 μM.

### Liposome Fusion Assays

Liposome fusion was assayed at 37°C or 24°C. Fusion reactions of 200 μl were assembled from three pre-mixes: two mixes of liposomes in RB150/Mg^2+^ and one mix of auxiliary factor GTP or their respective buffers. All components were incubated directly in 96-well plates, incubated for 30 or 60 min and content and/or lipid mixing signals were recorded at intervals of 60 s in a Infinite200 PRO (Tecan) fluorescent plate reader (MB::NBD-FRET Ex: 460 nm; Em: 538 nm; PhycoE::Cy5-FRET, Ex: 496nm; Em: 670 nm; MB dequenching, Ex: 370 nm; Em: 465 nm). For content mixing and lipid dequenching reactions, maximal values were determined after addition of 0.1% (wt/vol) thesit to the samples. FRET efficiency *E* defined as 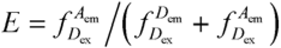 where 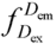 represents the donor emission intensity after donor excitation and 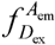 the acceptor emission intensity after donor excitation. For combination analysis of lipid dequenching and content mixing, excitation wavelength of PhycoE::Cy5-FRET changed to 535 nm. Liposome tethering was observed at 24 °C by sample turbidity changes at 350 nm with 0.5 μM protein and 1 mg ml^−1^ liposomes.

### Fluorescence Microscopy

For visualizing labeled vesicles, a Delta Vision Elite (GE Healthcare, Applied Precision) equipped with an Insight SSI^™^ illumination, a CoolSnap HQ2 CCD camera was used. Images were taken with a 100× oil PSF U-Plan S-Apo 1.4 NA objective. A four color standard set InsightSSI unit with following excitation wavelengths (DAPI 390/18 nm, Green 475/28, Orange 542/27, Cy5 632/22 nm); single band pass emission wavelengths (DAPI 435/48 nm, Green 573/36, Orange 594/45, Cy5 679/34 nm) and a suitable polychroic for DAPI/Green/Orange/Cy5 were used. Analysis of the images was performed using ImageJ. All imaging experiments were performed several times with biological replicates.

## Acknowledgements

The authors acknowledge funding from China Scholarship Council (CSC, a fellowship to L.G) and the Deutsche Forschungsgemeinschaft (DFG, INST 86/1452-1 and BR2915/4.1 to M.B.).

